# Stable Polymer Bilayers for Protein Channel Recordings at High Guanidinium Chloride Concentrations

**DOI:** 10.1101/2020.12.29.424722

**Authors:** Luning Yu, Xinqi Kang, Mohammad Amin Alibakhshi, Mikhail Pavlenok, Michael Niederweis, Meni Wanunu

## Abstract

Use of chaotropic reagents is common in biophysical characterization of biomolecules. When the study involves transmembrane protein channels, the stability of the protein channel and supporting bilayer membrane must be considered. In this letter we show that planar bilayers composed of poly(1,2-butadiene)-b-poly(ethylene oxide) diblock copolymer are stable and leak-free at high guanidinium chloride concentrations, in contrast to diphytanoyl phosphatidylcholine bilayers which exhibit deleterious leakage under similar conditions. Further, insertion and functional analysis of channels such as α-hemolysin and MspA are straightforward in these polymer membranes. Finally, we demonstrate that α-hemolysin channels maintain their structural integrity at 2M guanidinium chloride concentrations using blunt DNA hairpins as molecular reporters.

---

Planar bilayer membranes are commonly used to probe isolated trans-membrane channels using a patch-clamp setup(1). In this setup, two electrodes are used to apply voltage to electrolyte solutions present across a bilayer membrane, which drives ion flow across any available path through the membrane, for example, a trans-membrane protein channel such as α-hemolysin from *Staphylococcus aureus* (**Figure 1A**). Typically, the lipid bilayer is impermeable to ions, which allows sensitive measurements of ion flux across the membrane protein. However, in some conditions the electrolyte contains agents that compromise the bilayer membrane stability, which makes single-channel measurements impossible to conduct. Diphytanoyl phosphatidylcholine (DPhPC) is a common lipid often used to make bilayer membranes for single-channel measurements of different analytes. However, at high concentrations of guanidinium chloride (GdmCl), a common reagent that induces protein unfolding, DPhPC membranes exhibit significant permeability, and experiments are limited to low voltage and moderate GdmCl concentrations (2–5).

**Figure 1.**
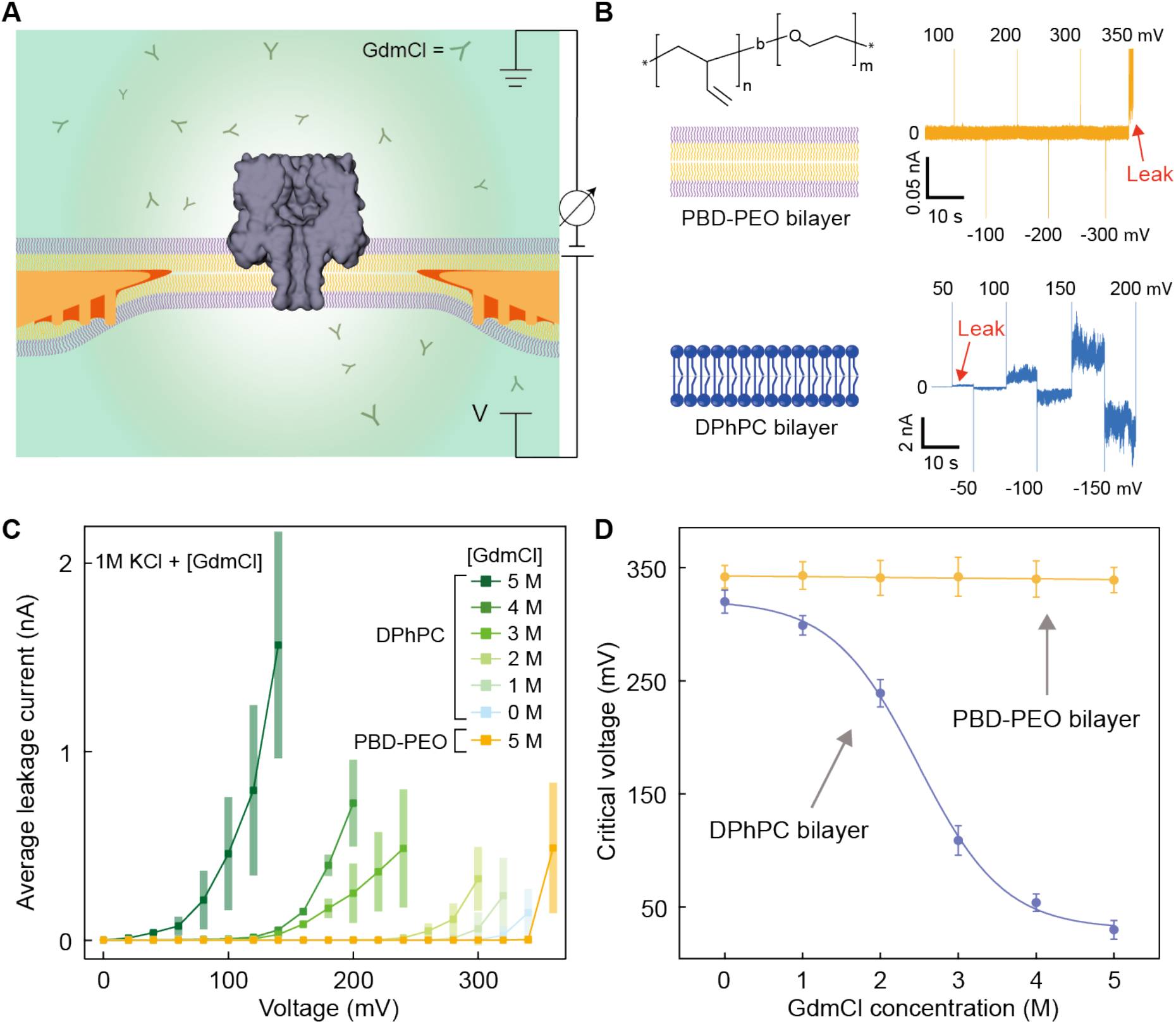
Permeability of DPhPC bilayers and PBD-PEO bilayers in high guanidinium chloride (GdmCl) concentrations. **A)** Cartoon side-view of a bilayer membrane supported by a wedge-on-pillar aperture(9). A single α-hemolysin channel inserted through the bilayer is shown. A patch-clamp amplifier is used to record ion flux through the bilayer and trans-membrane channel for electrolytes containing GdmCl. **B)** Current vs. time traces for PBD-PEO bilayers (chemical structure shown) and DPhPC bilayers when voltage increments are applied in opposite polarities in order to test permeability (electrolyte: 5 M GdmCl, 1 M KCl, 10 mM Tris, pH 7.5). Non-zero current fluctuations correspond to ion leakage through the membranes (indicated by red arrows). Traces were low-pass filtered to 10 kHz using the Axopatch internal filter and digitally sampled at 16-bit, 250 kHz using a DAQ card (National Instruments). **C)** Average leakage current vs. voltage for DPhPC and PBD-PEO bilayer membranes at indicated GdmCl concentrations (+1 M KCl). Error bars represent the standard deviation based on five sets of measurements. **D)** Critical voltage vs. [GdmCl] for PBD-PEO (yellow) and DPhPC (blue) bilayer membranes, determined by the voltage at which mean leakage current exceeds 5 pA. Line and sigmoidal curves are guides to the eye.

In search of a suitable replacement we have tested the amphiphilic block copolymer poly(1,2-butadiene)-b-poly(ethylene oxide) (abbreviated PBD-PEO, cat # P41807C-BdEO, Polymer Source, Montreal, Quebec, Canada). **Figure 1B** shows the chemical structure of PBD_n_-PEO_m_ (*n* ~ 11, *m* ~ 8), which has an estimated core bilayer thickness of ~4.4 nm (see Supporting Material (SM), **Figure S1**)(6), similar to commonly used DPhPC bilayers (~5.0 nm)(7, 8) which we use to benchmark against here (cat # 850356C, Avanti Polar Lipids, Alabaster, AL). After pipette-painting thin organic membrane solutions in decane across our 100 μm diameter wedge-on-pillar(9) aperture, we gradually thin the membrane by passing air bubbles across it until a true bilayer is formed, indicated by a rise in membrane capacitance to 70±10 pF, and also by the observation of spontaneous insertion of protein channels after their addition to the top (*cis*) chamber. Also shown in **Figure 1B** are excerpts of current vs. time recordings that probe ion permeability for 1 M KCl + 5M GdmCl buffer, obtained by applying alternating polarity DC voltages with increasing magnitudes and monitoring current as a function of time using a patch-clamp amplifier (Axopatch 200B, Molecular Devices, San Jose, CA). Leakage current vs. voltage statistics for different electrolytes are summarized in **Figure 1C**, where the error bars represent standard deviations of the leakage currents. To obtain these data we applied sequential 20 mV increments (positive polarity only) and monitored the current leakage. Even in 5 M GdmCl, the PBD-PEO membrane is impermeable up to 340 mV, whereas significant ion permeability was observed for DPhPC membranes at lower voltages for all electrolyte conditions (for representative raw traces see SM, **Figure S2**). In **Figure 1D** we present the critical voltage as a function of GdmCl concentration for DPhPC and PBD-PEO membranes, defined as the voltage at which the average DC current (I_avg_) exceeds 5 pA. The rapid decline in voltage stability for DPhPC suggests a significant membrane destabilization, possibly through formation of interactions with phosphate groups of DPhPC(10, 11). In contrast, PBD-PEO has an outstanding chemical resilience at GdmCl concentrations sufficient to denature most proteins.

In **Figure 2A** we present current traces before and immediately after single-channel insertion into PBD-PEO bilayer membranes, for both α-hemolysin from *S. aureus* (Sigma-Aldrich, Cat # H9395) and an engineered MspA channel from *Mycobacterium smegmatis*, M2-MspA(12). As seen in the traces, both M2-MspA and α-hemolysin produce stable current levels, similar to ones that are routinely observed with DPhPC membranes (see power spectra in SM, **Figure S3**). In **Figure 2B**, a 2.5 s current recording at 150 mV of a single M2-MspA channel is shown with spikes caused by the addition of a short (15-nt) single-stranded DNA to the *cis* chamber (for an extended trace see SM, **Figure S4**). The channel shows normal activity and a high rate of events caused by DNA interactions with the M2-MspA channel. In **Figure 2C**, a 52 s current recording at 100 mV of a α-hemolysin channel is shown with spikes caused by the addition of a 60-nt single-stranded DNA to the *cis* chamber (for an extended trace see SM, **Figure S5**). Similar to M2-MspA, the α-hemolysin channel is stable and active, as evidenced by the stable baseline current and a “healthy” rate of events. Based on these data we confirm the suitability of PBD-PEO bilayer membranes for single-channel recordings. Finally, in **Figure 2D** we present histograms of experimental bilayer membrane lifetimes, measured by monitoring the current as a function of time for a sealed fluidic cell and recording the time it takes for the bilayers to rupture in 1M KCl, 10mM Tris, pH 7.5 electrolyte. The mean lifetime of PBD-PEO bilayers is 2-fold to 3-fold higher than DPhPC membranes. It should be noted that we have used the PBD-PEO polymers (stored in chloroform at 4°C) for up to 17 months and have not observed any performance degradation, in contrast to DPhPC lipids, which typically degrade within months if not stored with care at −20°C under conditions that minimize oxidation/hydrolysis(13).

**Figure 2.**
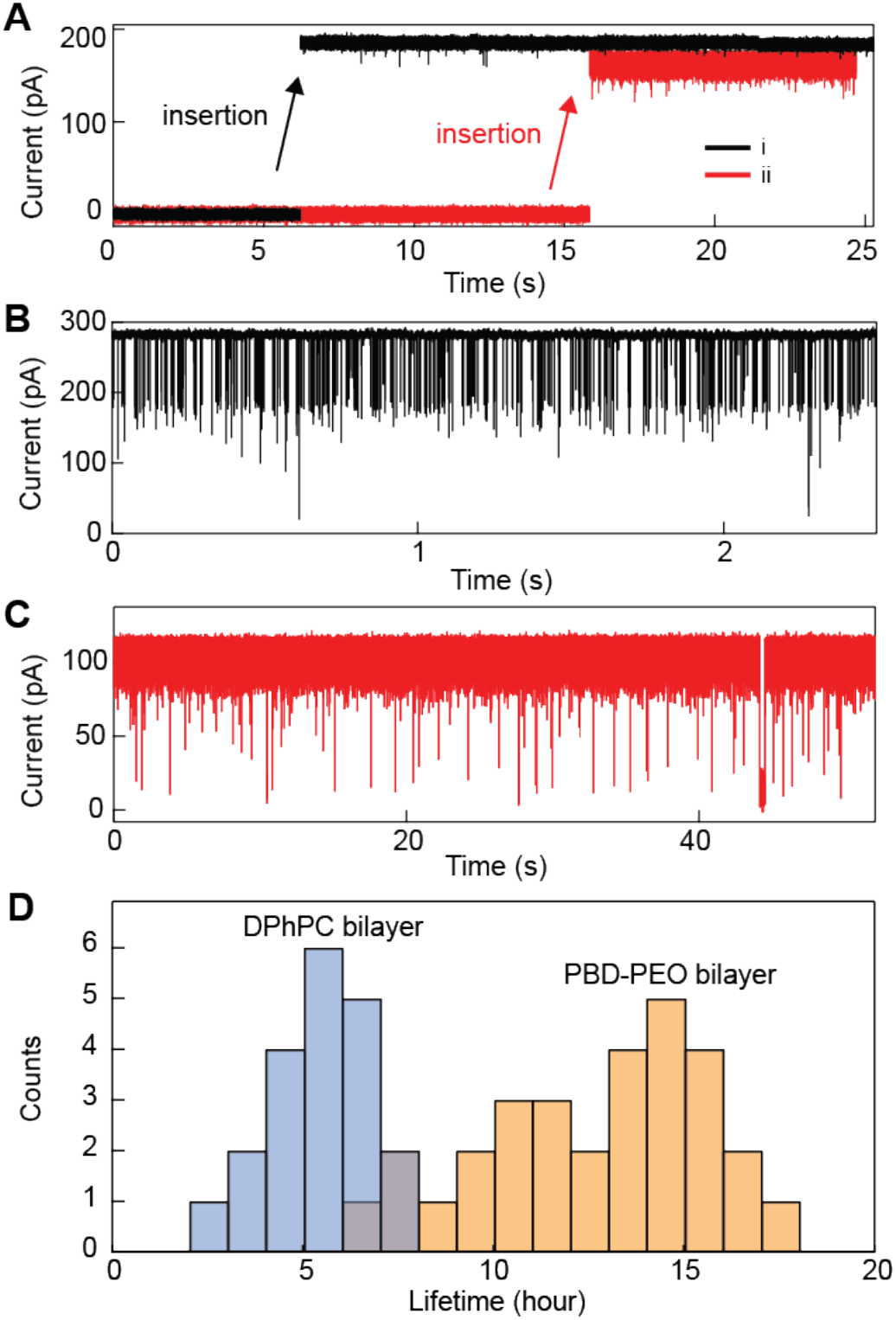
Single-channel measurements in PBD-PEO bilayer membranes. **A)** Current vs. time trace before and after single-channel insertion of i) M2-MspA (<I> = 184.9 pA) and ii) α-hemolysin (<I> = 164.5 pA) channels (electrolyte: 1 M KCl, 1.5 M GdmCl, 10 mM Tris, pH 7.5; V = 100 mV). **B)** Current recording of a M2-MspA channel when 1.8 μM 15-nt single-stranded DNA was placed in the *cis* chamber (electrolyte: 1 M KCl, 10 mM Tris, pH 7.5; V = 150 mV). **C)** Current recording of a α-hemolysin channel when 1 μM 60-nt single-stranded DNA was placed in the *cis* chamber (electrolyte: 1 M KCl, 10 mM Tris, pH 7.5; V = 100 mV). Traces for **B** and **C** were low-pass filtered to 10 kHz using the Axopatch internal filter and digitally sampled at 16-bit, 250 kHz using a DAQ card (National Instruments). **D)** Lifetime histograms measured for DPhPC (*n* = 20) and PBD-PEO (*n* = 30) bilayer membranes (electrolyte: 1 M KCl, 10 mM Tris, pH 7.5; V = 0).

Finally, we utilize the stable PBD-PEO bilayer membranes to test the structural integrity of the α–hemolysin pore at high GdmCl concentrations. It has been suggested that high urea concentrations might denature the α–hemolysin vestibule cap(14), although this has been questioned in subsequent measurements(15). Here we use DNA-based molecular probes to test the vestibule cap intactness. Short, blunt DNA hairpins are known to produce distinct signatures in α–hemolysin through long-lived interactions with the vestibule regions(16). These interactions are so intricate that DNA hairpins of different stem lengths can be distinguished with 1 bp resolution. In **Figure 3A**, we show a trace for a mixture of 4 bp and 6 bp blunt hairpins with sequences 5’-CTCGTTTTCGAG-3’ and 5’-CTAACGTTTTCGTTAG-3’, respectively, added to the *cis* chamber (Integrated DNA Technologies, Inc., Coralville, Iowa). Long-lived events with two characteristic amplitude levels were obtained, which correspond to the expected levels for these hairpin lengths(9, 16). For the same hairpin mixture in 2 M GdmCl + 1 M KCl buffer (**Figure 3B**), we find much faster events that exclusively contain two levels, an intermediate blockade level with relatively long (~ms) timescale, followed by a rapid deep level, which we attribute to translocation of the denatured DNA hairpin (for longer traces see SM, **Figure S6**). **Figure 3C** shows a scatter plot of fractional blockades vs. event dwell times for the two experiments in **A** and **B**. The scatter plots reveal dwell times that are on average 2-3 orders of magnitude shorter than for 1 M KCl buffer alone, suggesting that 2 M GdmCl affects hairpin stability. The effect of urea on disrupting DNA and RNA secondary structures are well known and have been reported to facilitate transport of structured nucleic acids through α–hemolysin channels(15). Despite the impact on DNA hairpin stability, the fractional blockade levels for the observed events are very similar (<2% difference) for both electrolyte conditions, which rules out appreciable structural changes to the α–hemolysin vestibule cap in high GdmCl concentrations.

**Figure 3.**
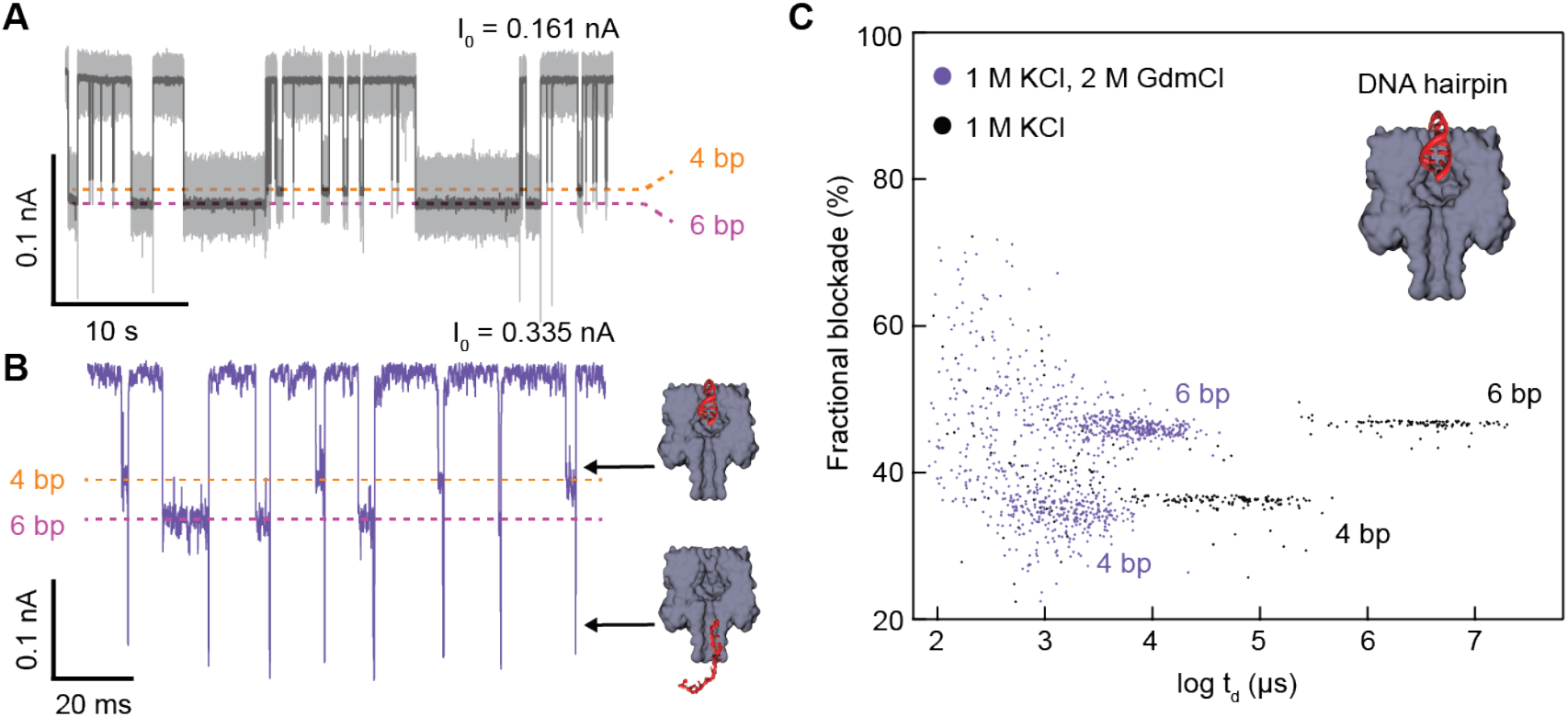
Single-channel measurements in PBD-PEO bilayer membranes. **A-B)** Current vs. time trace obtained for an α-hemolysin channel when a mixture of 4 bp (0.4 μM) and 6 bp (0.6 μM) blunt DNA hairpins with a T4 loop is added to the *cis* chamber containing 1M KCl (**A**) and 2 M GdmCl + 1 M KCl (**B**) (electrolyte also contained 10 mM Tris, pH 7.5; V = 175 mV) To the right of the trace in **B**, we show a cartoon of proposed states that correspond to the current signals obtained during 4-bp hairpin occlusion of the vestibule and unzipped hairpin translocation, respectively. **C)**Scatter plots of fractional current blockades vs. dwell times for the indicated experiments in **A**, **B**.

In summary, we have shown that bilayer membranes composed of PBD-PEO amphiphilic di-block copolymer are suitable for single-channel recordings in presence of high GdmCl concentrations. The lifetime of PBD-PEO bilayers is superior to that of DPhPC lipid membranes, and reconstitution of protein channels readily occurs. We have utilized this finding to show that the vestibule cap of α-hemolysin maintains its integrity at 2 M GdmCl concentrations, while significantly destabilizing the secondary duplex structures of short blunt DNA hairpins.

## Acknowledgment

This work was supported by the National Institutes of Health (NIH) grant R21HG010543 to M.N.

## Supporting Material

Supporting traces and extended raw data.

