## Supporting Material for "Stable Polymer Bilayers for Protein Channel Recordings at High Guanidinium Chloride Concentrations"

**Table of Contents:**

1) **Figure S1:** Poly-butadiene (PBD) core thickness vs. number of monomers in a PBD-PEO block copolymer bilayer.

2) **Figure S2**: Representative current vs. time traces for studying the voltage stability of DPhPC and PBD-PEO bilayer membranes (supplements Figure 1D).

3) **Figure S3**: Power spectral density plots of single-channel M2-MspA and $\alpha$-hemolysin (supplements Figure 2A).

4) **Figure S4**: Extended recording of M2-MspA channel in presence of short (15-nt) single-stranded DNA (supplements Figure 2B).

5) **Figure S5**: Extended recording of $\alpha$-hemolysin channel in presence of 60-nt single-stranded DNA (supplements Figure 2C).

6) **Figure S6**: Extended recordings for the DNA hairpin mixture data (supplements Figure 3A, 3B).

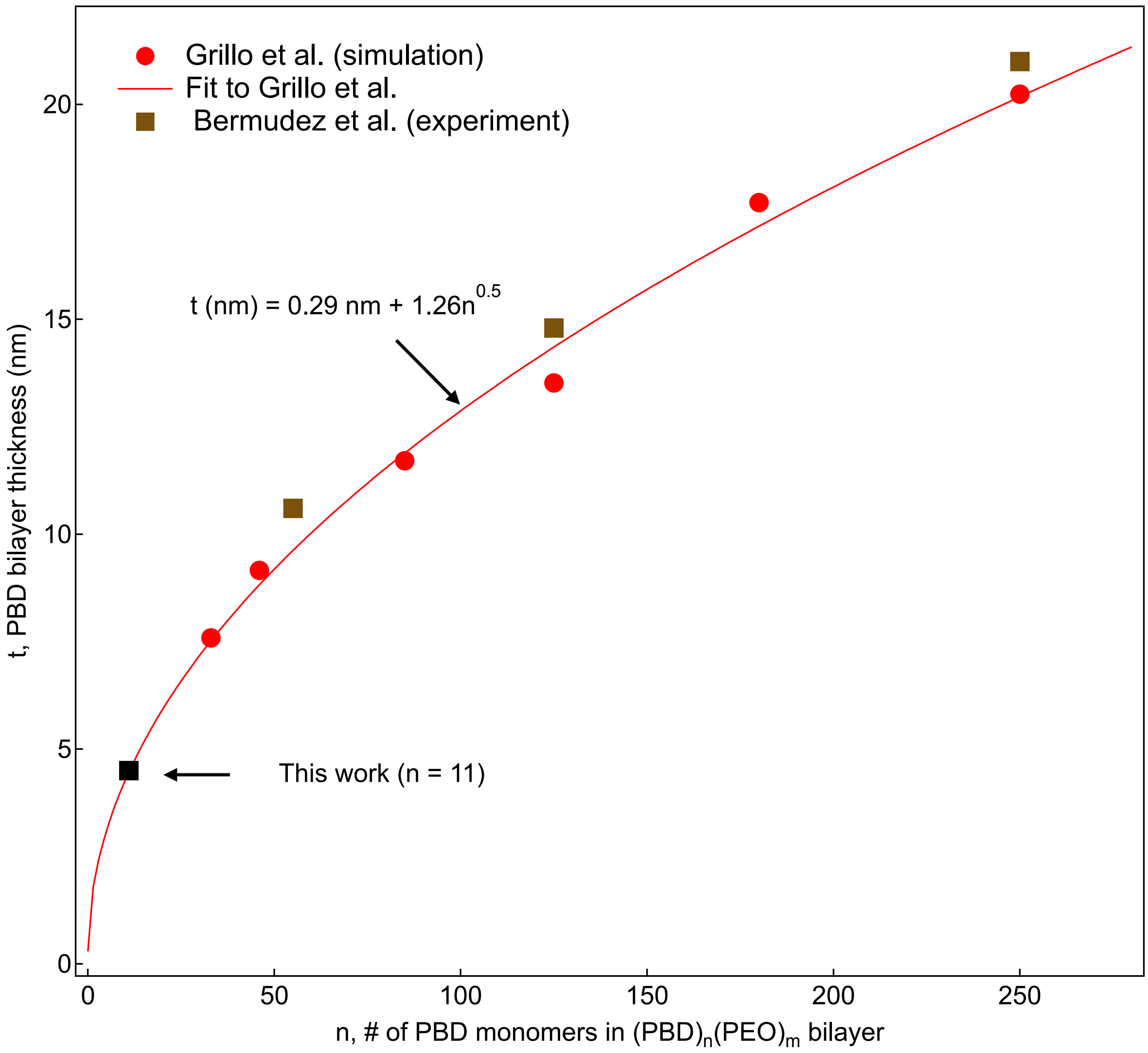


**Figure S1.** Estimated core thickness of the poly-butadiene (PBD) bilayer vs. number of PBD monomers in a PBD-PEO block copolymer. Red circles are from Grillo et al. (1), and brown squares are from Bermudez et al. (2), Table III. The red curve shows a power-law fit (exponent 0.5) to the data, from which our polymer bilayer's core thickness was estimated to be 4.4 nm.

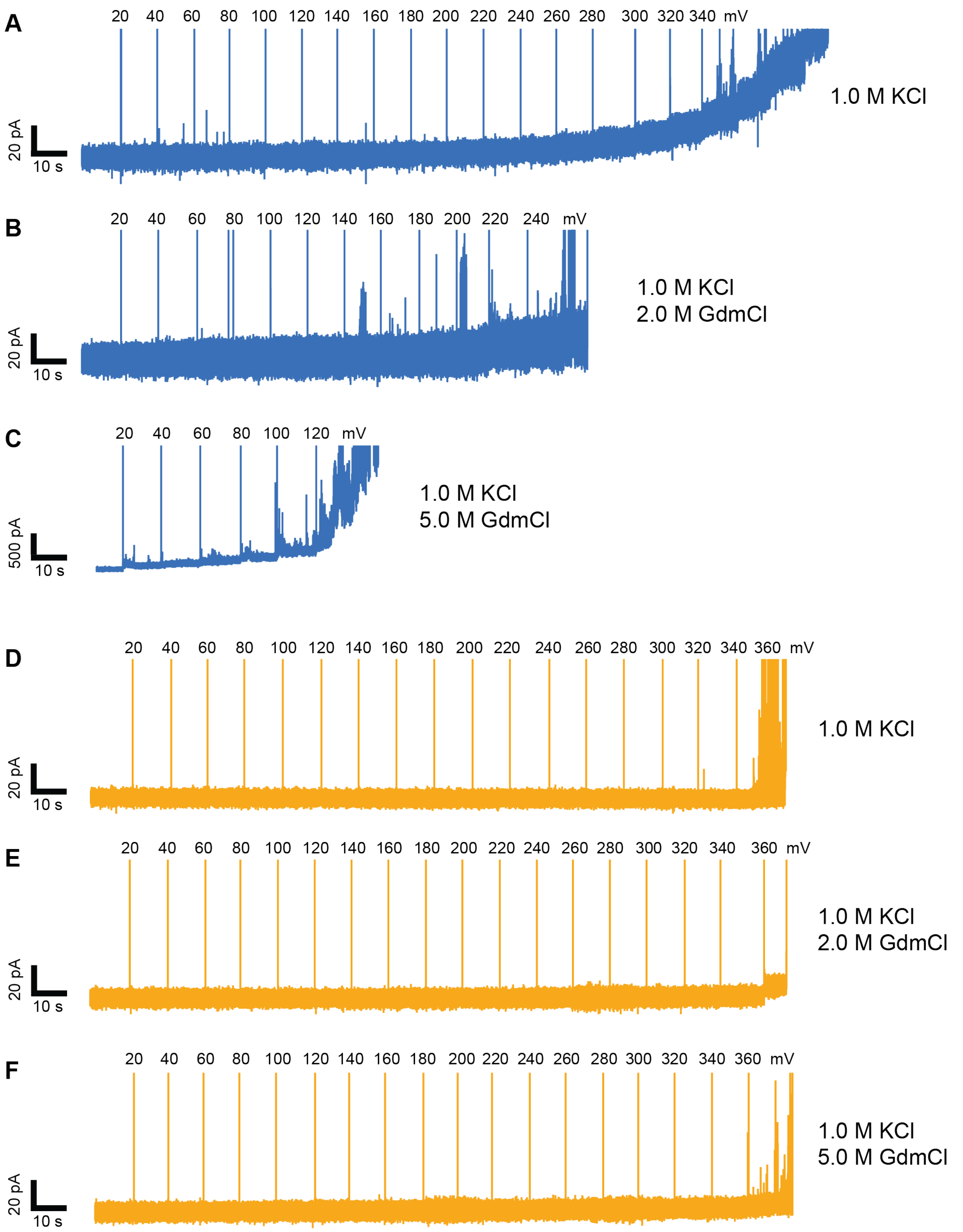


**Figure S2.** Representative current traces obtained for voltage stability tests of (**A-C**) DPhPC and (**D-F**) PBD-PEO bilayer membranes (used to generate the plot in Figure 1D).

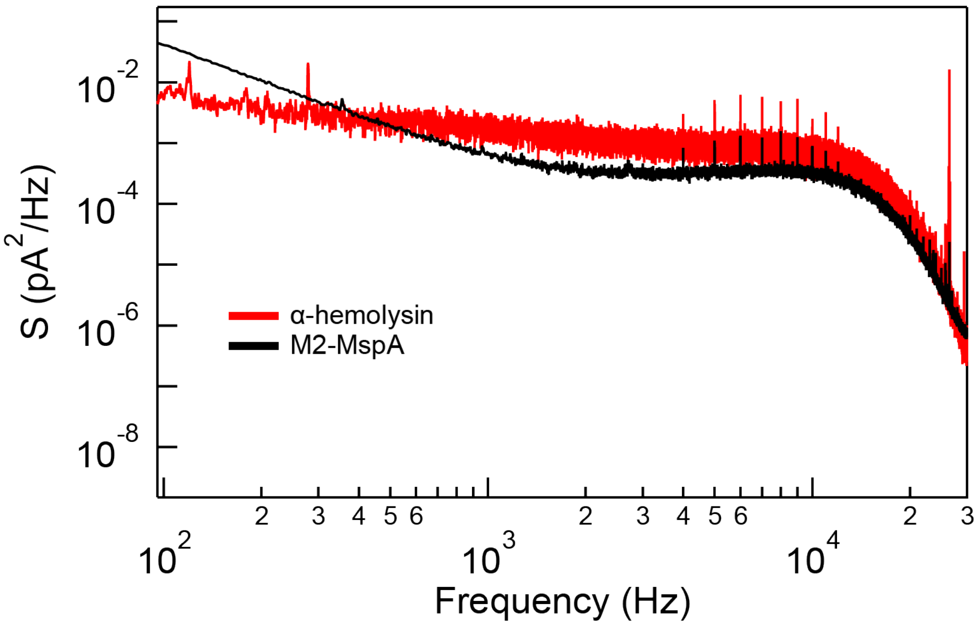


**Figure S3.** Power spectral density plots of the traces shown in Figure 2A (M2-MspA and α-hemolysin channels).

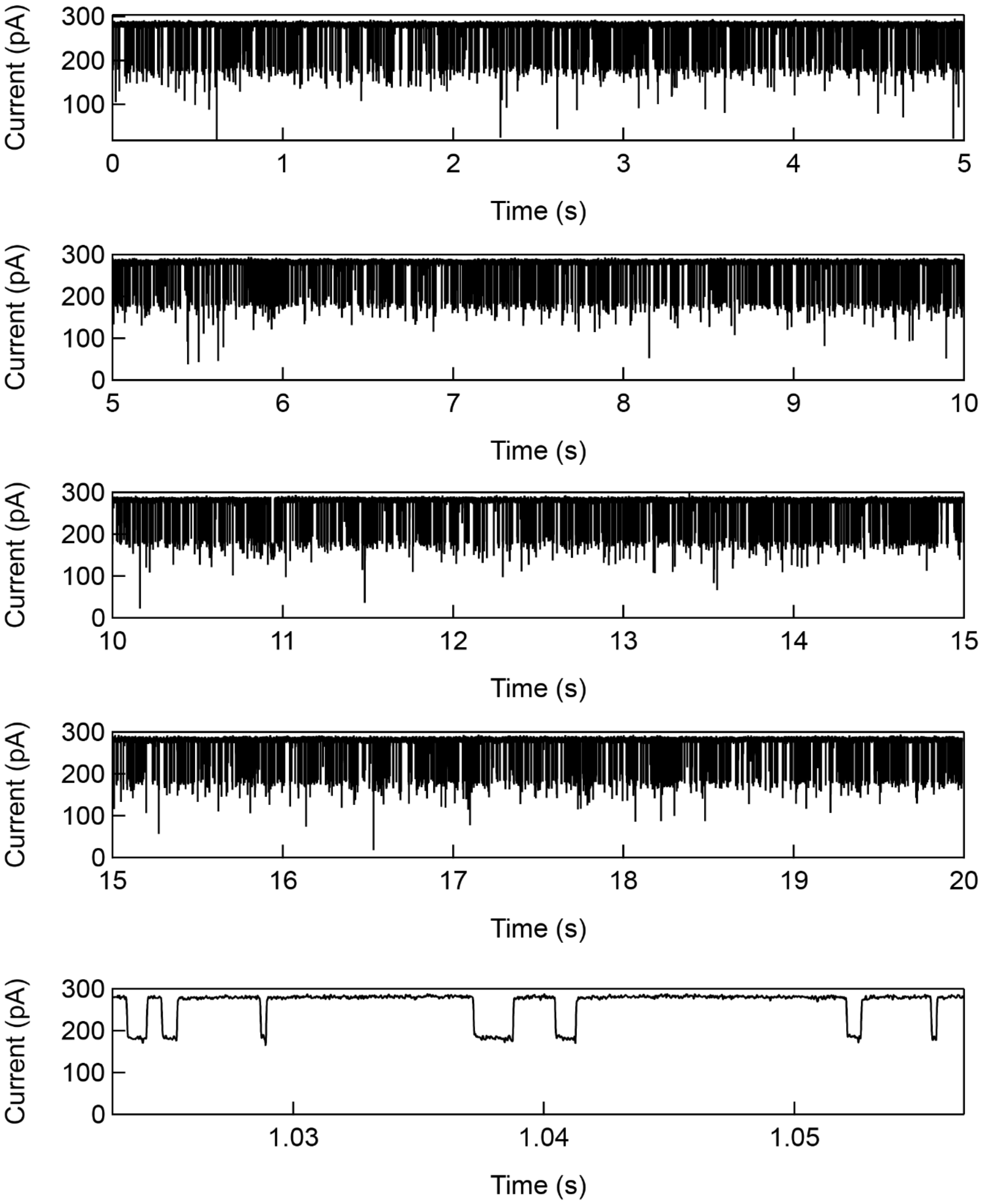


**Figure S4.** Extended recording of M2-MspA channel in presence of short (15-nt) single-stranded DNA (sequence: 5'-CTG CTT GCA TCG TAG-3').

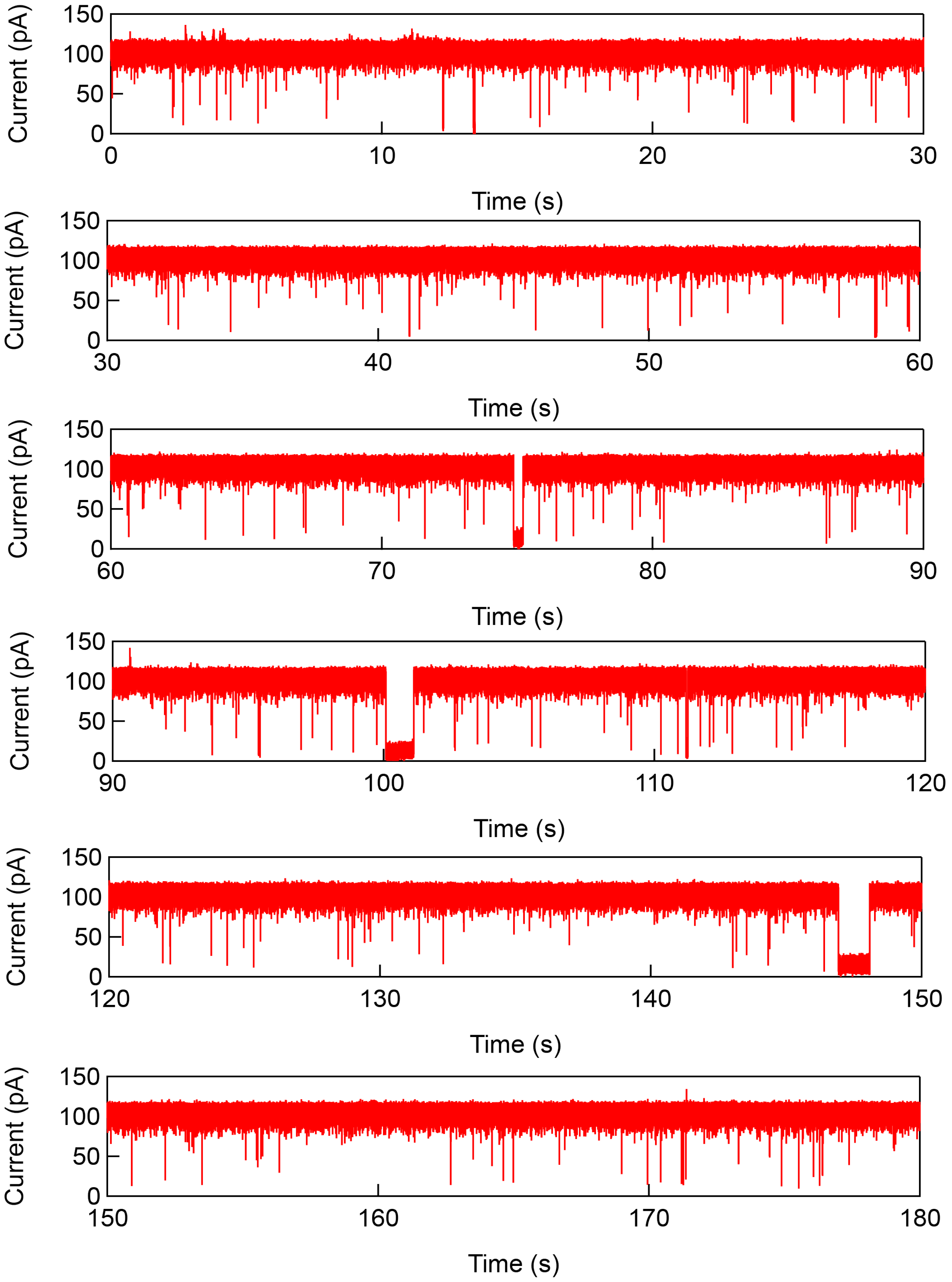


**Figure S5.** Extended recording of α-hemolysin channel in presence of short (60-nt) single-stranded DNA (sequence: 5'-AAAAAAAAACCCCCCCCCCCTTGATGCACTGGCTTAAAAAAAAAACCCCCCCCCCCCCC-3').

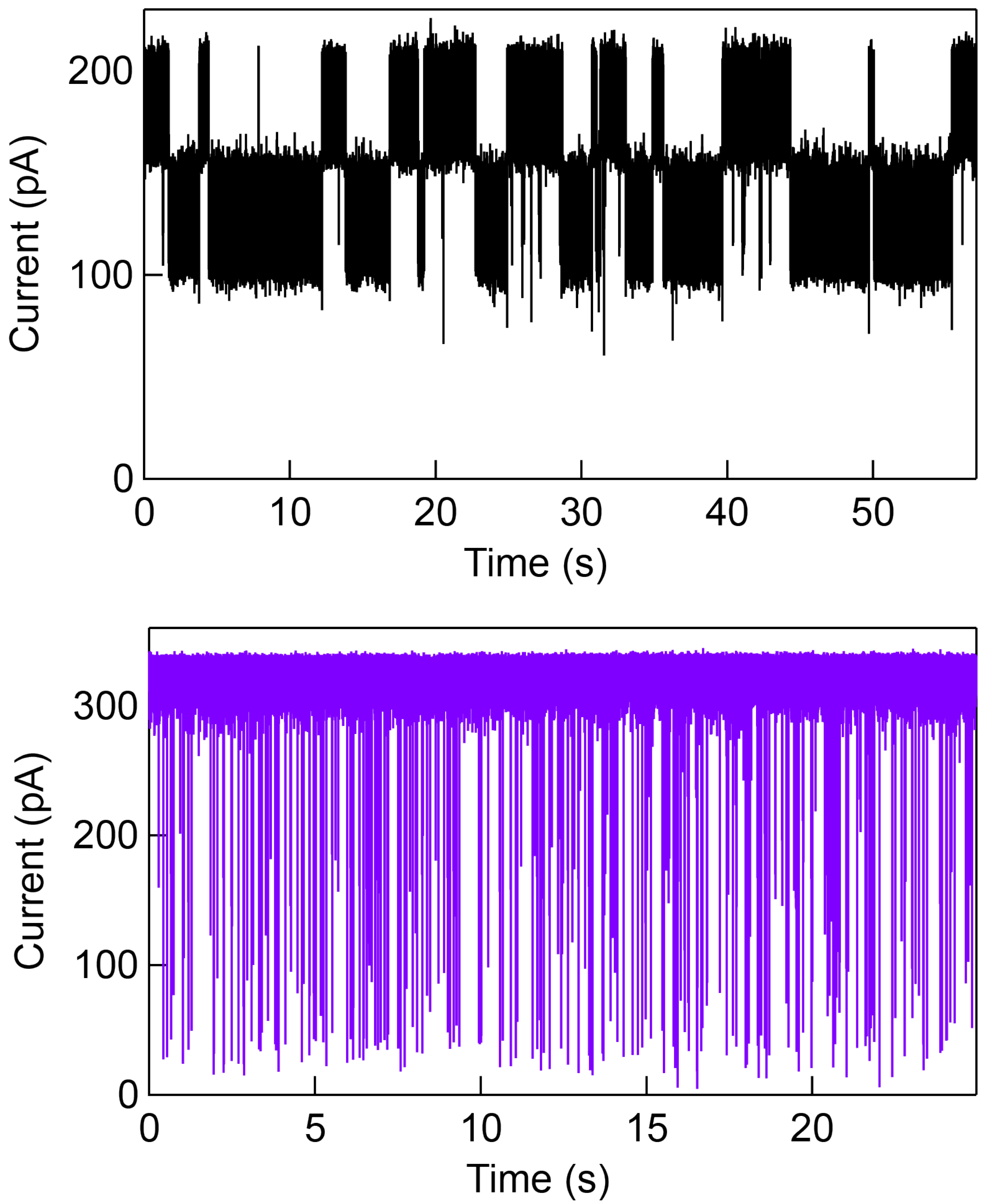


**Figure S6.** Current vs. time recordings for a 4:6 mixture of 4 bp and 6 bp DNA hairpins added to the *cis* chamber for a α–hemolysin channel in 1M KCl, 10mM Tris, pH 7.5, V = 175 mV (top), and in 2M GdmCl (1M KCl, 10mM Tris, pH 7.5, V = 175 mV (bottom). In 1 M KCl, the hairpin stability is high enough that dwell times are orders of magnitude longer than for the case of 2 M GdmCl, in which case hairpin melting occurs readily (above traces supplement the data in Figure 3 of the main text).
